# Phase variation in LPS O-antigen biosynthetic gene cluster of the rice pathogen *Xanthomonas oryzae* pv. *oryzae*

**DOI:** 10.1101/2023.04.27.538605

**Authors:** Vishnu Narayanan Madhavan, Kanika Bansal, Sanjeet Kumar, Samriti Midha, Prabhu B. Patil, Hitendra Kumar Patel, Ramesh V. Sonti

**Author notes:** **Author for correspondence:** Ramesh V. Sonti, ICGEB Campus, Aruna Asaf Ali Marg, 110067, New Delhi, India.

## Abstract

Bacteria respond to environmental cues in different ways. Phase variation is one such adaptation where heritable and reversible changes in DNA aid bacteria to alter the expression of specific genes. The bacterial plant pathogen *Xanthomonas oryzae* pv. *oryzae* (Xoo) causes the serious bacterial blight disease of rice. The mucoid phenotype of Xoo colonies is attributed to the secreted exopolysaccharide (EPS), xanthan gum. Spontaneous non-mucoid variants of Xoo which are deficient in EPS production and virulence were observed to accumulate in long-term stationary phase cultures. This phenomenon was termed stationary phase variation and variant colonies as stationary phase variants (SPV). Several but not all of these SPVs have been earlier described to carry spontaneous insertions of endogenous insertion sequence elements in the *gum* operon which encodes genes involved in EPS biosynthesis. In this study, we show that a number of SPVs harbour variations in the lipopolysaccharide (LPS) outer antigen (O-antigen) biosynthetic gene cluster. The data revealed that the vast majority of variations are due to either insertion of endogenous insertion sequence (IS) elements or slipped strand mispairing (SSM). Also, it was observed that many of these SPVs exhibited reversion to wild type mucoid phenotype via restoration of the wild type genotype. The results indicate that the phenomenon of phase variation is occurring in the LPS O-antigen biosynthetic gene cluster of Xoo.

## Introduction

Phase and antigenic variation in bacteria is the alteration between two or more phenotypes in a heritable and reversible manner. Phase variation thus enables bacteria to maintain wild type and one or more phase variant populations and is thought to aid the bacteria in adapting to changes in their environment. Phase and antigenic variation have been extensively studied and established in free-living and animal pathogenic bacteria. A wide array of genes, encoding structural to biosynthetic proteins have been shown to undergo phase or antigenic variation. The mechanisms of phase and antigenic variation include slipped-strand mispairing (SSM), general recombination, conserved site-specific recombination, transposition of insertion sequence (IS) elements and DNA methylation (Woude and Bäumler, 2004).

Bacteria produce different kinds of polysaccharides including those linked to the membrane (like capsular polysaccharides) or secreted exopolysaccharides (EPS) as well as lipopolysaccharides (LPS) and lipo-oligosaccharides (LOS) that are structural components of the outer membrane in Gram-negative bacteria. The genes involved in the biosynthesis or modifications of the polysaccharide portion of LPS (known as outer antigen or O-antigen) and LOS have been shown to undergo phase variations via slippage of homo or heteropolymeric tract of nucleotides leading to insertions or deletions, a phenomenon referred to as slipped-strand mispairing (SSM) (Weiser, Love and Moxon, 1989; Appelmelk *et al*., 1999; Wang *et al*., 1999; Linton *et al*., 2000; Gilbert *et al*., 2002). The reversible variation of EPS biosynthetic gene, *epsG* give rise to a non-mucoid, crenated phenotype in *Pseudoaltremonas atlantica* via insertion and precise excision of IS element *IS492* (Bartlett, Wright and Silverman, 1988; Bartlett and Silverman, 1989; Higgins, Carpenter and Karls, 2007). The *icaC* gene of *icaABCD* operon involved in the biosynthesis of EPS known as polysaccharide intercellular adhesin (PIA) in *Staphylococcus aureus* and *Staphylococcus epidermidis* undergoes phase variation via transposition involving the *IS256* IS element (Ziebuhr *et al*., 1999; Kiem *et al*., 2004). Further, PIA phase variation through SSM in *icaC* gene has been described for *Staphylococcus aureus* (Brooks and Jefferson, 2014). Insertion of *IS5*-like IS element insertions in the capsular polysaccharide operon has been attributed to the translucent phenotype of *Pseudoaltremonas lipolytia* (Zeng *et al*., 2019).

Like other members of the bacterial genus *Xanthomonas*, the rice pathogen *Xanthomonas oryzae* pv. *oryzae* (Xoo) has a characteristic mucoid phenotype due to copious secretion of an EPS called xanthan gum or xanthan. Non-mucoid colonies deficient in EPS and virulence were observed to accumulate spontaneously in long-term stationary phase cultures of Xoo. These strains were termed stationary phase variants (SPVs) (Rajeshwari and Sonti, 1997). The non-mucoid phenotype of four of ten such SPVs was complemented by a cosmid clone (pSD1) carrying the EPS biosynthetic gene cluster that is referred as the *gum* operon. All these four SPVs were shown to harbour insertions of endogenous IS elements in the *gumM* gene of the *gum* operon. Three of these strains had insertions of the IS element *ISXo1* and one strain had an insertion of the *ISXo2* IS element. The nature of variations in the remaining SPVs was not characterised (Rajeshwari and Sonti, 2000).

In the present study, we have isolated a fresh set of SPVs using the previously established method (Rajeshwari and Sonti, 1997). Most of the SPVs isolated harbour variations in the LPS O-antigen biosynthetic gene cluster that give rise to the non-mucoid phenotype. Characterisation of these SPVs indicated that the variations include endogenous IS element insertions and SSM. Additionally, a number of these SPVs exhibited reversion to wild type like mucoid phenotype. Upon investigation of such ‘reverted’ colonies it was revealed that the sequence of the LPS O-antigen biosynthetic cluster at the site of variation has been restored to that of the wild type. These results suggest that phase variation at the LPS O-antigen biosynthetic gene cluster reversibly affects the production of LPS O-antigen and EPS, as well as the virulence of Xoo.

## Results

### The non-mucoid phenotype of SPVs is complemented by a cosmid clone containing the LPS O-antigen biosynthetic gene cluster

SPVs were isolated from 14 independently grown long-term stationary phase cultures of wild type Xoo strain BXO43 using the methodology described previously (Rajeshwari and Sonti, 1997). The SPVs were identified based on their non-mucoid colony morphology, which is indicative of an EPS deficient phenotype (Figures 1A and 1B). Previous studies have indicated that mutations in either the *gum* operon or the LPS O-antigen biosynthetic gene cluster of Xoo result in EPS deficiency and exhibit a non-mucoid colony phenotype (Dharmapuri *et al*., 2001). Hence the first question addressed was whether the 14 SPVs harbour variations in either the LPS O-antigen biosynthetic gene cluster or the *gum* operon. To test this, complementation analysis was performed by independentyly transforming the 14 SPVs with pSD1 and pSD5 cosmid clones which contain *gum* operon and LPS O-antigen biosynthetic gene cluster respectively (Dharmapuri and Sonti, 1999; Dharmapuri *et al*., 2001). The restoration of wild type mucoid phenotype was scored visually. Among the 14 SPVs, the phenotype of 13 SPVs was restored to mucoid phenotype by pSD5 cosmid clone (Figures 1C, 1D, and Table 1). Further, these 13 SPVs lacked the characteristic O-antigen containing band in their LPS profile (Figure 2). Taken together, the results indicate that the non-mucoid phenotype of these 13 SPVs is due to mutations in the LPS O-antigen biosynthetic gene cluster. The phenotype of one SPV strain, SPV15 was complemented by the pSD1 cosmid clone (containing the *gum* operon) indicating that this strain is mutated in the *gum* operon. As expected, the LPS profile of this SPV strain was similar to that of the wild type (Figure 2).

**Figure 1.**
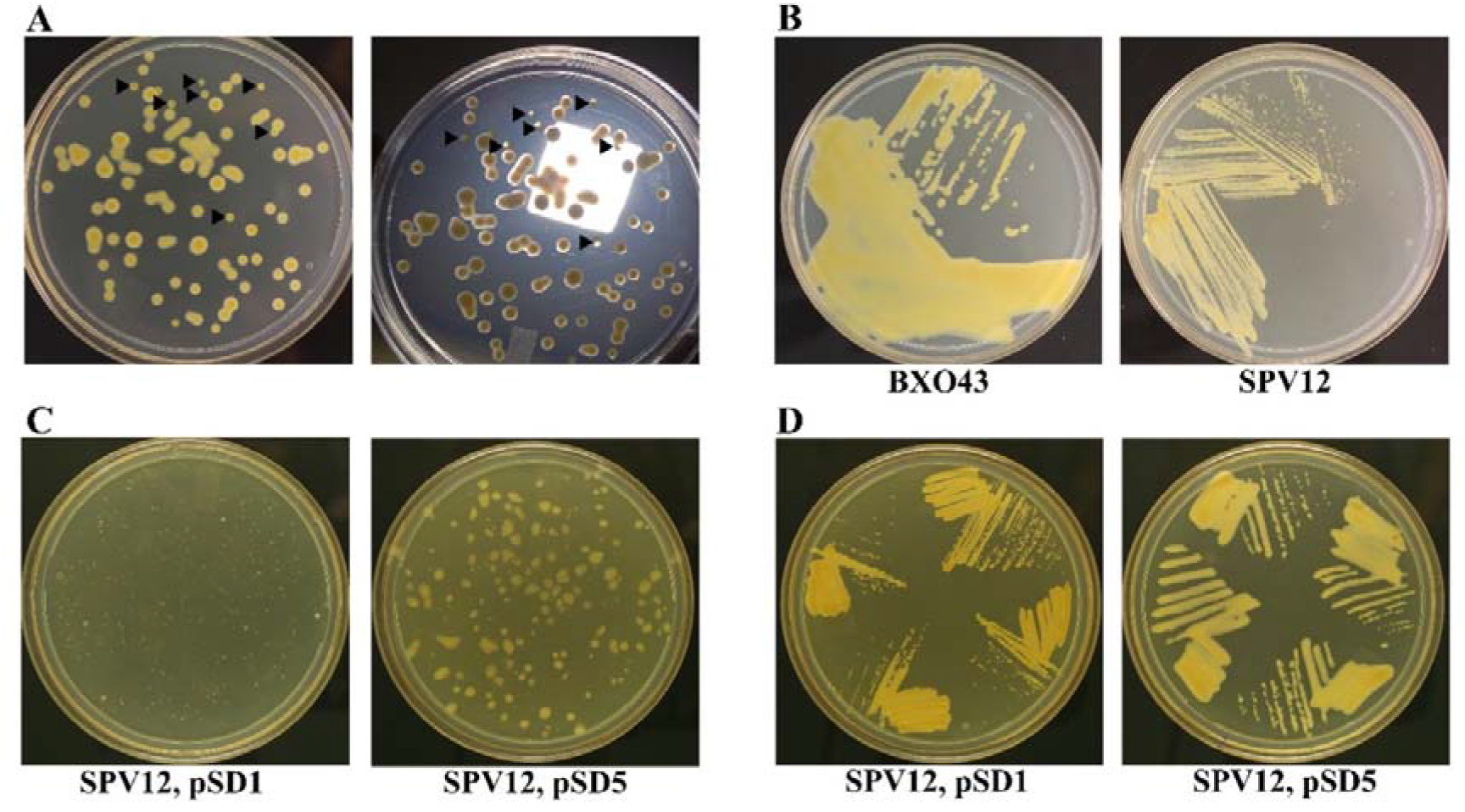
Non-mucoid phenotype of SPV and phenotype complementation. **(A)** Representative image of a dilution plate from day 8 of a prolonged stationary phase culture of wild type Xoo, BXO43. The arrows indicate the small, EPS deficient SPV colonies (left) which appear translucent under direct light (right) compared to opaque, large, mucoid wild type colonies. **(B)** Wild type Xoo BXO43 and SPV12 streaked on plates for comparison of phenotype. **(C, D)** Representative images of SPV12 transformed with either *gum* locus (pSD1) or LPS O-antigen biosynthetic cluster (pSD5) containing cosmids. **(C)** Individual colonies on selection media and **(D)** colonies streaked on plates to observe the phenotype. In this example, non-mucoid phenotype of SPV12 is complemented by pSD5.

**Table 1.** Complementation analysis of the non-mucoid phenotype of the 14 SPVs using genomic DNA library clones.

| Cosmid | Cluster | SPVs in which mucoid phenotype is restored |
| --- | --- | --- |
| pSD1 | EPS biosynthetic cluster (*gum* operon) | 15 |
| pSD5 | LPS O-antigen biosynthetic cluster | 1, 2, 3, 4, 5, 11, 12, 16, 22, 26, 31, 32, 33 |

**Figure 2.**
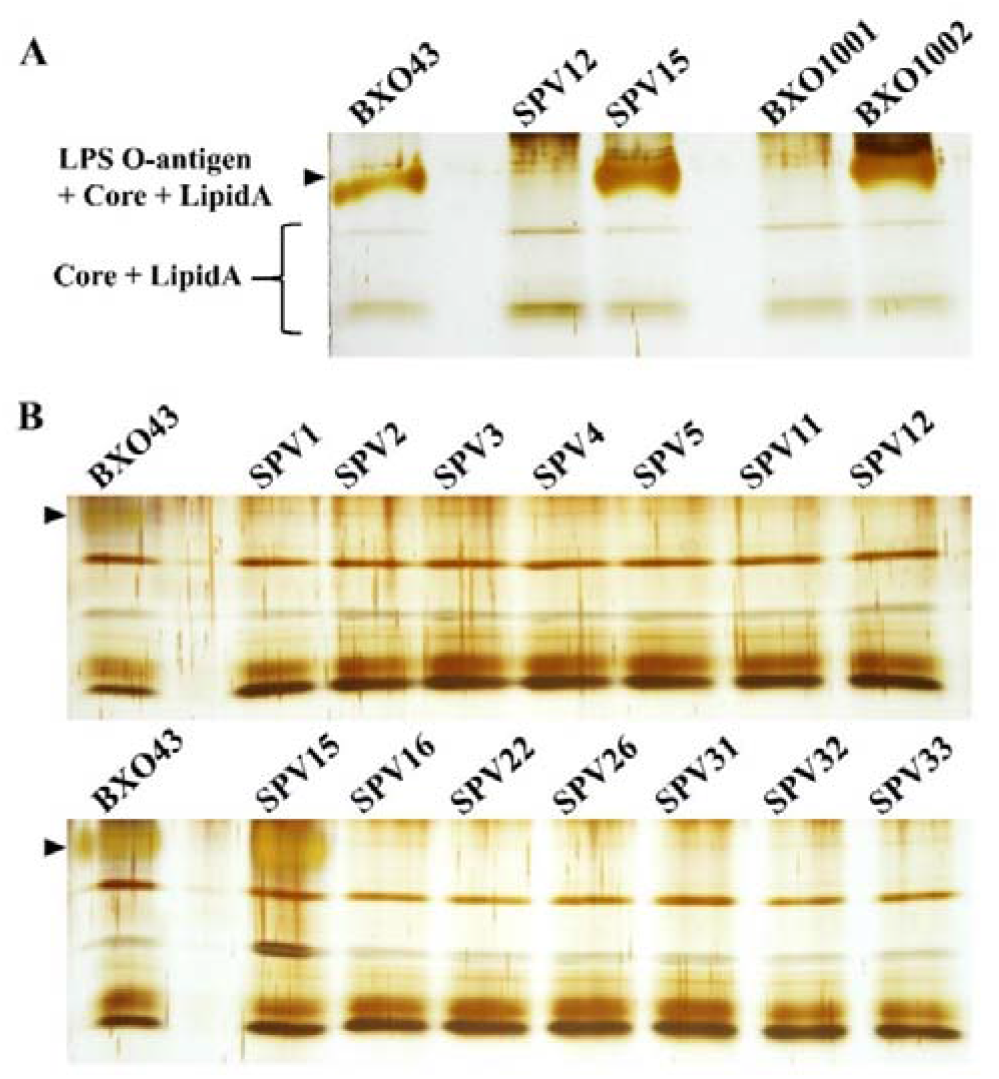
LPS profile of SPVs. **(A)** Comparison of LPS isolated from wild type Xoo BXO43, SPVs 12 and 15 in which the non-mucoid phenotype can be complemented by pSD5 and pSD1 respectively with BXO1001 and BXO1002 which are characterised mutants of LPS O-antigen biosynthetic gene cluster and *gum* locus respectively. **(B)** Comparison of LPS profile of all 14 SPVs and wild type Xoo strain BXO43. The 13 SPVs in which the non-mucoid phenotype can be complemented by LPS O-antigen biosynthetic gene cluster containing cosmid (pSD5), lacks the characteristic band containing O-antigen. The black arrows indicate the characteristic band containing LPS O-antigen. The experiment was repeated three times and similar results were obtained.

### Some SPVs harbour insertions of endogenous IS elements in the LPS O-antigen biosynthetic gene cluster

The O-antigen biosynthetic gene cluster was PCR amplified using appropriate primers to generate 13 overlapping PCR amplicons of length ∼ 500 bp and overlap of ∼ 100 bp between adjacent amplicons. Individual primer pairs were designed to amplify the two transporter channel protein genes (Figure 3A). Analysis of the PCR products from 14 SPVs and the wild type strain using the aforementioned primers indicated that several SPVs carry approximately 1 kb insertions in the LPS O-antigen biosynthetic genes (Figure 3B). Analysis of the Whole Genome Shotgun (WGS) sequencing data of the SPVs that harboured insertions revealed that the corresponding genes contain contig breaks. Additionally, SPV15, the only SPV for which the pSD1 cosmid clone (containing the *gum* operon) restored the mucoid phenotype was observed to have a contig break in the *gumB* gene. The PCR products from the SPVs that appear to carry insertions were sequenced and were indeed found to contain insertions of endogenous IS elements. The results revealed that 6 SPVs, *viz*. SPV 1, 11, 12, 16, 22, and 26 harbour IS element insertions in the LPS O-antigen biosynthetic gene cluster and SPV15 harboured an IS element insertion in the *gumB* gene (Figure 3C). Among these seven SPVs, four SPVs have insertions of *ISXo1* and target site duplication (TSD) of 4 base pairs. Eight base pair TSD was observed for insertions of *ISXo2* in two SPVs and insertion of *ISXoo13* in one SPV (Table 2).

**Table 2.** Characteristics of IS element insertions in SPVs.

| SPV | Location of insertion (Gene, bp) | IS element | Orientation of transposase gene with respect to the native locus gene | Target site duplication sequence |
| --- | --- | --- | --- | --- |
| 22 | *wxoA*, 740bp | *ISXoo13* | Opposite | AATTTATG |
| 11 | *wxoB*, 1714bp | *ISXo1* | Opposite | TTAG |
| 12 | *wxoB*, 1714bp | *ISXo1* | Same | TTAG |
| 1 | *wxoC*, 129bp | *ISXo1* | Same | TTAG |
| 26 | *wxoC*, 252bp | *ISXo1* | Opposite | CTAG |
| 16 | *wxoC*, 529bp | *ISXo2* | Same | TTTCTTTT |
| 15 | *gumB*, 489bp | *ISXo2* | Opposite | AAGATGAT |

**Figure 3.**
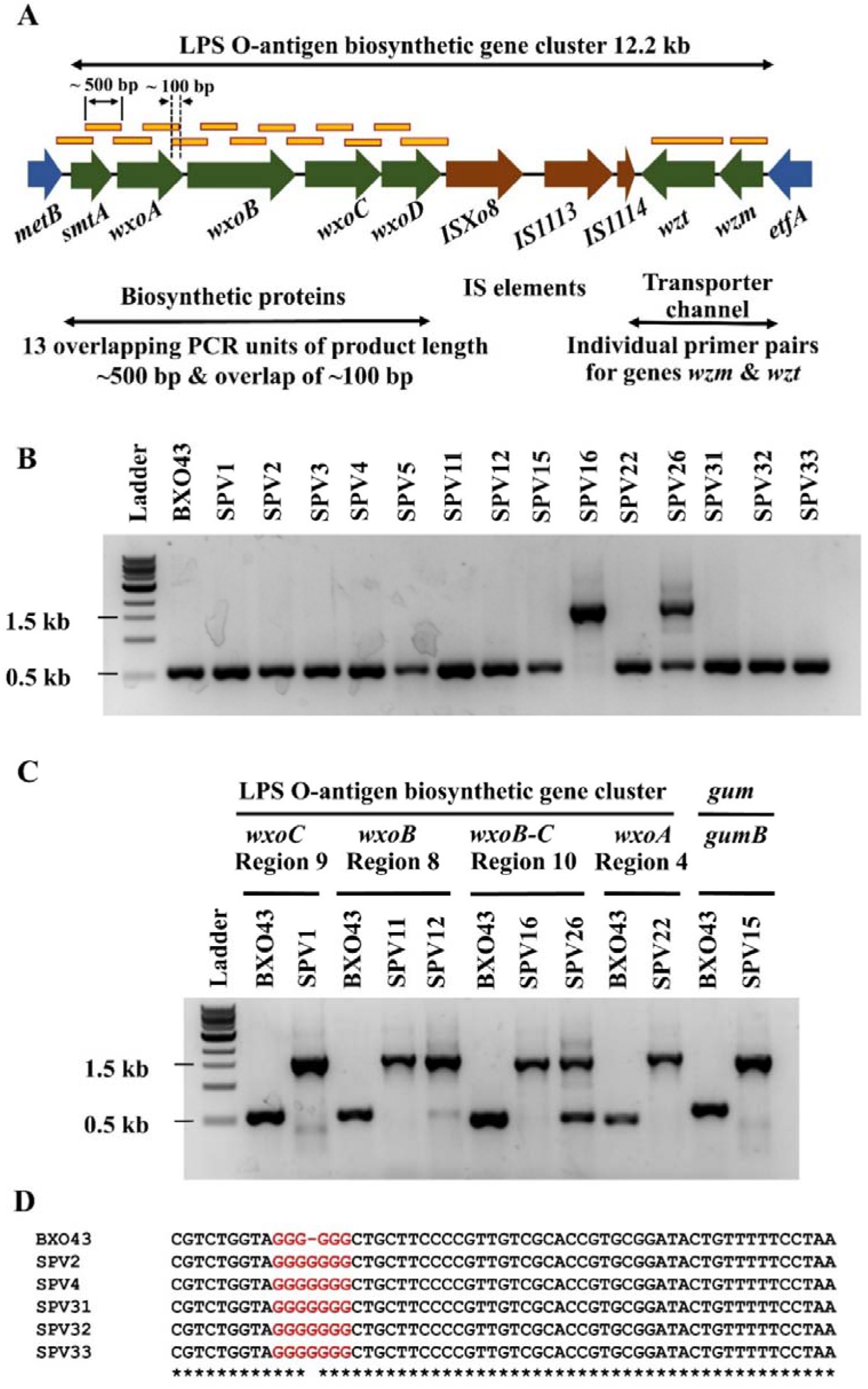
Characterisation of variations in SPVs. **(A)** Schematic of LPS O-antigen biosynthetic cluster (green arrows), the cluster location is conserved in *Xanthomonas* between the flanking genes *metB* and *etfA* (blue arrows), and the IS elements within the cluster (brown arrows) (Patil, Bogdanove and Sonti, 2007). The overlapping regions PCR amplified for screening variations are represented in yellow rectangles. **(B)** Representative agarose gel image of insertion screening. The PCR products for the ‘Region10’ were amplified using genomic DNA isolated from BXO43 and all 14 SPVs as templates. The ∼ 1.5 kb bands in SPV 16 and 26 are indicative of insertions. **(C)** Agarose gel image of the SPVs which showed insertions. Presence of wild type like ∼ 500 bp band in SPV26 is indicative reversion. **(D)** Multiple sequence alignment snippet of the portion containing the G-hexamer of *wxoA* gene. PCR products for the Region 4 to 5 were amplified using genomic DNA isolated from five SPVs and BXO43 were sequenced.

### Slipped strand mispairing (SSM) at a G-hexamer in the *wxoA* gene of the LPS O-antigen biosynthetic gene cluster

The WGS sequence data were examined to understand the nature of variation in the remaining 7 SPVs for which the phenotype did not appear to be due to IS element. The data revealed that five SPVs, *viz*. SPV 2, 4, 31, 32, and 33 have a G-heptamer instead of a G-hexamer in the *wxoA* gene of the LPS O-antigen biosynthetic gene cluster. This result was confirmed by PCR amplification and sequencing of the corresponding region from the *wxoA* gene (Figure 3D). The SSM leads to altered open reading frame of *wxoA* gene and also generate a premature termination codon. The insertion of an extra nucleotide at a run of nucleotide repeats suggests slipped strand mispairing (SSM) at the G-hexamer of the *wxoA* gene.

### Genetic changes in SPV3 and SPV5

The SPV3 and SPV5 strains did not have any change in the G-hexamer of *wxoA* gene and we could not identify any specific IS element insertions. An examination of the WGS sequence data revealed that the SPV3 strain contains a premature stop codon in the *wxoA* gene. The SPV5 strain has a contig break in the *wxoA* gene which could not be amplified irrespective of multiple attempts using different polymerases and primer combinations.

### SPVs show phenotype and genotype reversion

During phenotyping, several but not all SPV strains were observed to revert to the wild type-phenotype and were classified into three categories. ‘Reverting type’ SPVs produced mucoid colonies or mucoid sectors amongst colonies dilution plated from exponential phase cultures. Reverting type SPVs also generated multiple mucoid sectors or complete mucoid phenotype when grown by spotting exponential phase cultures on plates. The frequency of reversion was observed to be 0.01-5% in exponential phase cultures. ‘Slow-reverting type’ SPVs produced mucoid regions occasionally and only when spotted on to plates and ‘non-reverting type’ SPVs did not generate any reversion phenotype in any assay conditions including serial passaging of exponential phase cultures. (Figures 4A, 4B, and Table 3).

**Table 3.** The reversion phenotypes observed in 14 SPVs.

| SPV strains | Phenotype |
| --- | --- |
| 11, 12, 16, 26, 31, 32, 33 | Reverting |
| 2, 4 | Slow reverting |
| 1, 3, 5, 15, 22 | Non-reverting |

**Figure 4.**
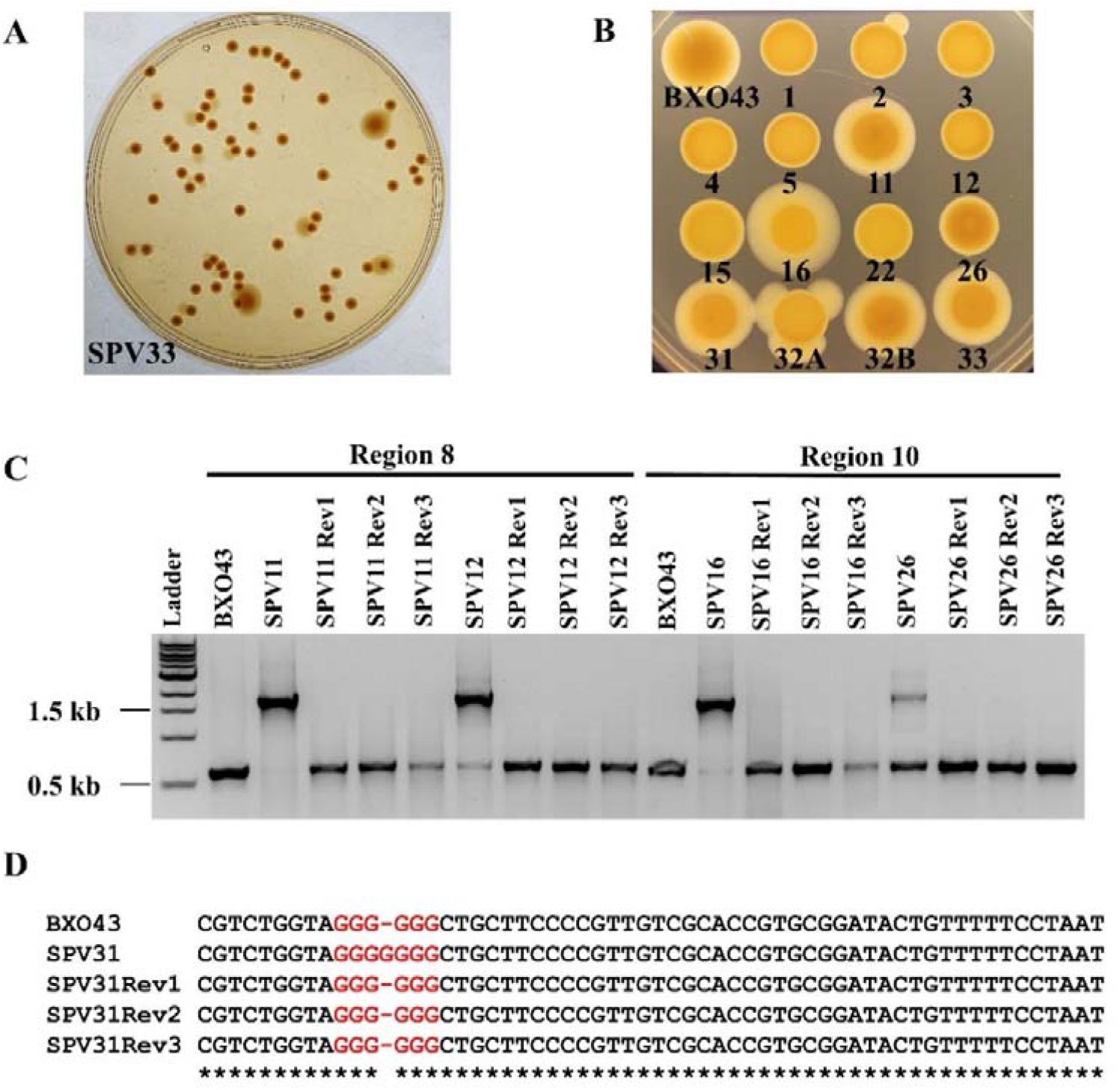
SPVs exhibit true reversion. **(A)** Dilution plating of SPV33 showing the reverting-type phenotype. The large dark opaque colonies arise from the reversion event that happened in the culture. Colonies with the white sectors indicate reversion events that happened after plating. **(B)** Representative image of wild type BXO43 and all 14 SPVs spotted on a PS agar plate. Reverting-type SPVs show multiple sectors, large halo or large mucoid region like wild type (SPV11, 12, 16, 26, 31, 32 and 33). Slow reverting-type SPVs show few sectors, at low frequency. SPV2 in the figure is an example here for slow reverting type (also SPV4 which is not showing sector in this particular set). The experiment was repeated four times and obtained similar results. **(C)** Agarose gel image for PCR products for the ‘regions’ that harbor IS element insertion with corresponding PCR primers and genomic DNA isolated from wild type BXO43, SPVs, and three independent isolates of reverted colonies from the respective SPV. Presence of the wild type like 500 bp bands in the reverted colonies indicates the restoration of wild type genotype. The wild type like bands in SPV lanes are indicative of revertants in the broth culture. **(D)** Representative image showing restoration of G-hexamer of *wxoA* in reverted colonies isolated from SPVs which exhibited G-heptamer. Snippet from multiple sequence alignment for the sequence obtained for PCR product of the region 3 containing the G-hexamer (indicated in red colour) of *wxoA*. Genomic DNA was isolated from wild type BXO43, SPV31 and three independent reverted colonies from SPV31. Similar results were obtained for all SPVs that exhibited SSM at *wxoA, viz*.SPV 2, 4, 32 and 33.

To understand the nature of this phenotypic reversion three independent ‘revertants’ (colonies with mucoid phenotype) were isolated from each of the SPVs and genotyped. The revertants obtained from SPVs 11, 12, 16, and 26 appeared to have lost the IS elements as PCR analysis of the target region resulted in PCR amplified products that were approximately 500 bp in size, similar to that of the wild type Xoo strain (Figure 4C). The sequencing of these PCR amplified products revealed that the inserted IS elements have undergone a precise excision event restoring the wild-type genotype.

All the five SPVs that were identified to have SSM at G-hexamer of *wxoA viz*. SPV 2, 4, 31, 32, and 33 also showed the reversion phenotypes. Sequencing of the target region of the *wxoA* gene revealed that the revertants have a G-hexamer like the wild type strain (Figure 4D). These results conclude that the phenotypic reversion of non-mucoid SPVs to wild type mucoid phenotype is due to the restoration of wild type genotype i.e. the SPVs exhibit true reversion.

### Reverting type SPVs successfully generate virulence lesion upon *in-planta* inoculation

Xoo strains with mutations in either the EPS or the LPS O-antigen biosynthetic gene cluster are virulence deficient (Dharmapuri and Sonti, 1999; Dharmapuri *et al*., 2001). The virulence phenotype of SPVs was assessed by clip inoculation of rice leaves. Lesions were observed in leaves inoculated with the reverting type SPVs while leaves inoculated with either the slow or non-reverting SPVs did not produce significant lesions (Figure 5). The bacteria were isolated from the leaves showing virulence lesions and were observed to have mucoid colonies. This suggests that restoration of wild type genotype and phenotype in reverting type SPVs also restores the capability to cause disease.

**Figure 5.**
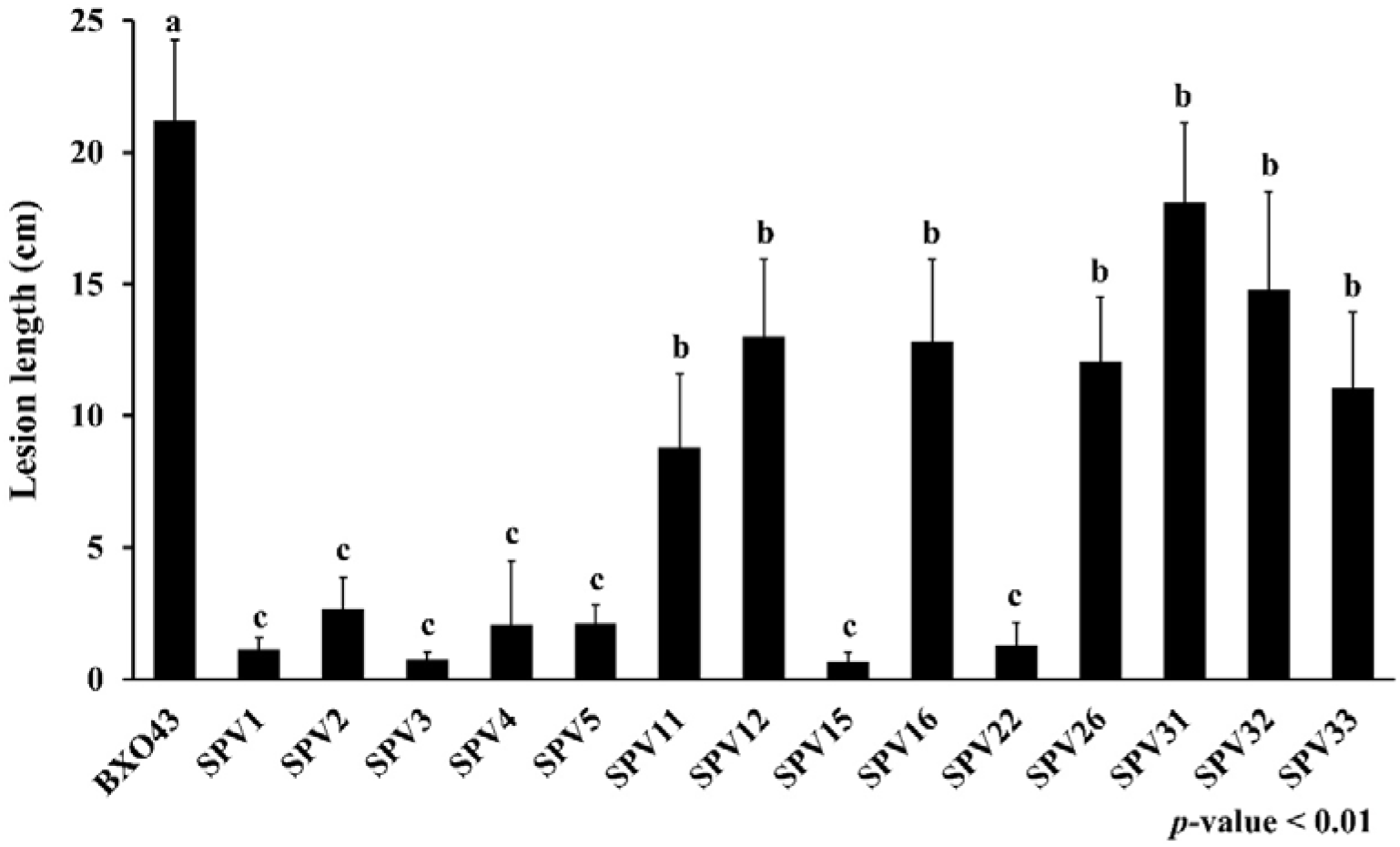
Bar graph showing the average lesion lengths caused by all 14 SPVs and wild type BXO43. The Xoo strains were clip inoculated on 60-80 days old TN1 rice leaves and lesions were measured 15 days post inoculation. Error bars represent standard deviation calculated from 10 or more leaves. The different letter labelling on the bars (a, b, and c) indicate a significant difference in lesion length from others using the unpaired two-tailed Student’s *t-* test (*p*-value < 0.01). The experiment was repeated three times and similar results were obtained.

## Discussion

Phenotype switching with the evidence of reversion to wild type phenotype *in-vitro* or *in-vivo* has been studied extensively in animal pathogenic bacteria and has been referred to as phase variation. The mechanisms which bring about heritable but reversible changes in DNA have been suggested to aid bacteria in adapting to environmental conditions (Woude and Bäumler, 2004). Among plant pathogenic bacteria, non-pathogenic, EPS deficient variants isolated in still broth cultures of *Ralstonia solanacearum* were described to undergo phenotype reversion when inoculated on the host plant. IS element insertions and tandem duplications in the *phcA* gene (a transcription regulator) were the mechanisms involved in the characterised variants (Poussier *et al*., 2003). In the previous study from our lab, one of the SPV strain that had been isolated from the wild type Xoo strain was also observed to undergo phenotypic reversion phenotype upon *in-planta* inoculation (Rajeshwari and Sonti, 1997). Phenotype switching between swarming colonies with EPS deficient phenotype and wild type have been described in *Xanthomonas campestris* pv. *campestris* (Kamoun and Kado, 1990). However, the underlying mechanisms were unexplored.

In the present study, we have uncovered the mechanisms of the reversible phase variation occurring in Xoo during prolonged stationary phase growth. Results indicated that, besides the mutations in the *gum* operon, SPVs also harbour mutations in the LPS O-antigen biosynthetic genes cluster leading to EPS deficiency. Among the 14 SPVs characterised in the present study, 13 SPVs harbour mutations in the LPS O-antigen biosynthetic gene cluster and one SPV strain carried the mutation in the *gum* operon which are summarised along with reversion phenotypes in Table 4. Reversion phenotypes were observed in 9 of the 14 SPVs including all 5 SPVs that have SSM at the G-hexamer of *wxoA* exhibited. Among SPVs exhibiting SSM, SPVs 2 and 4 exhibit slow-reverting phenotype, possibly due to a second site variation. Out of the 7 SPVs harbouring IS element insertions, reversion could be detected in 4 of them. It is possible that the IS elements excise at a lower frequency in the remaining 3 SPVs because of which reversion could not be detected in our assay conditions. Insertion of *ISXo1* element was observed in a number of SPVs that were characterised in the present and our previous studies (R Rajeshwari and Sonti, 2000). One possibility is that this could be due to the small target site of 4 bp. Also, among the 4 SPVs harbouring *ISXo1* insertions, 3 showed reversion phenotypes. The target site of 4 bp might once again be a contributing factor, as it has a partial palindromic nature (CTAG or TTAG, i.e. YTAR). The high-frequency precision excision of IS elements in Xoo leaves us with questions that warrant future investigations. Do the transposases in IS elements have any growth-dependent expression or activity? Are there any additional trans-acting host factors involved in precise excision? Is the LPS O-antigen biosynthetic gene cluster a hotspot for transposition? One could begin addressing these questions by asking how the target sites, growth conditions or the host influence the excision events. To address this, the IS element insertions along with the target site duplications can be cloned into an antibiotic resistance gene, such that IS element excision would result in resistance to the antibiotic.

**Table 4.**
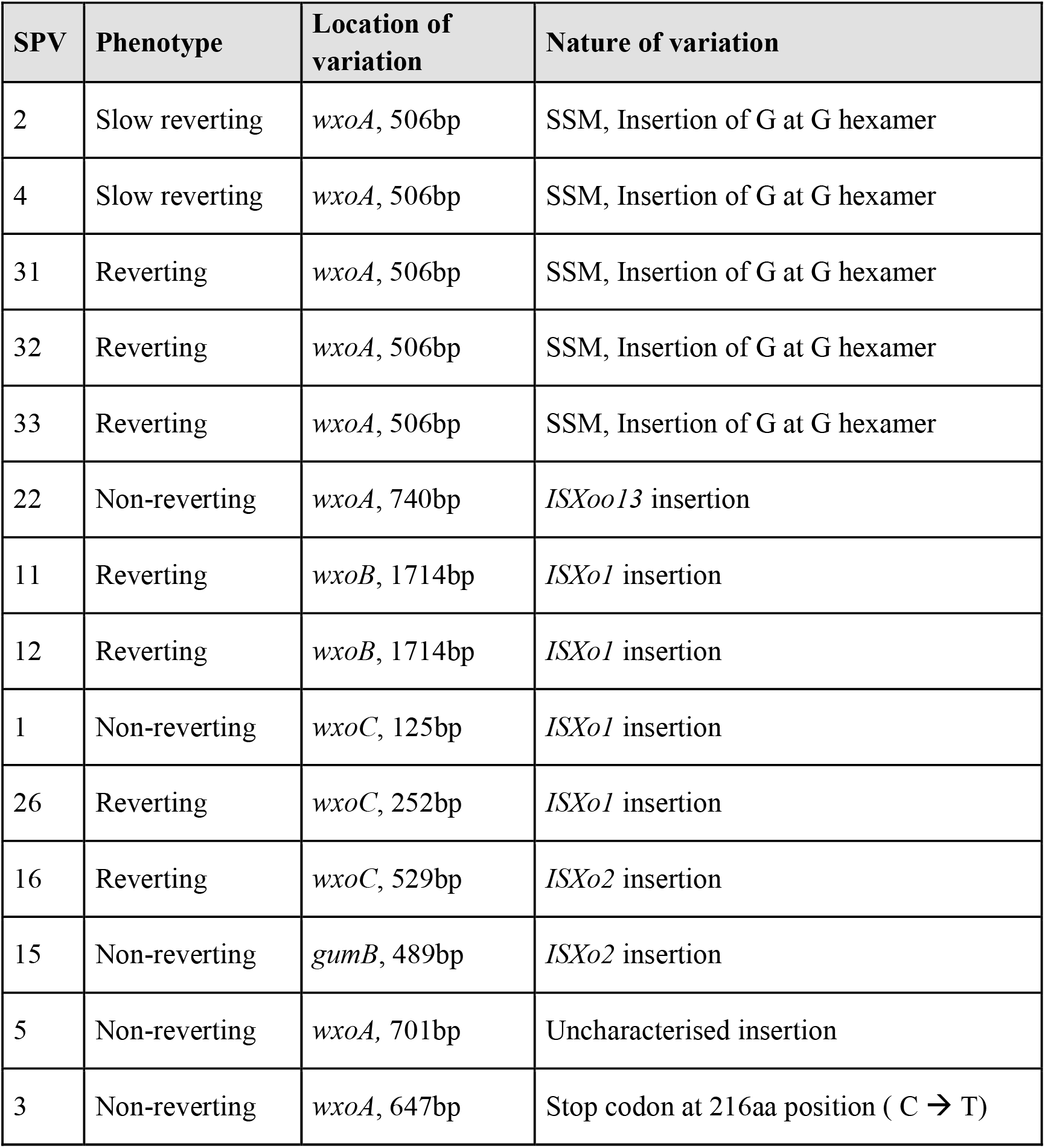
Nature and location of variations observed in all 14 SPVs.

Among the non-reverting type SPVs, WGS analysis revealed that SPV3 which harbours a stop codon in the *wxoA* gene contains more than 900 single nucleotide polymorphisms in its genome when compared to other SPVs and might be a possible mutator strain (Supplementary Table S1, S2). SPV5 in which the variation was located in *wxoA*, (contig break in WGS analysis) was not characterised since the region did not amplify even after multiple attempts. One possibility is that the region might be having a large insertion or have undergone a complex rearrangement following an IS element insertion.

Phase variation at the EPS biosynthetic locus has been described in other bacteria. Phase variation at the *icaC* gene involved in the production of polysaccharide intercellular adhesin (PIA) in *Staphylococcus aureus* has reduced biofilm formation, a critical requirement in its pathogenesis. However, the *icaC-*negative phase variant strains have been reported to have a survival advantage relative to wild type or complete *icaABCD* operon deletion, in co-culture experiments through an unknown mechanism (Brooks and Jefferson, 2014). Phase variation at the *epsG* gene of *P. atlantica* leads to a non-mucoid phenotype and reduced biofilm formation and it has been suggested that the EPS deficient phenotype aids in the dispersal of bacteria (Bartlett, Wright and Silverman, 1988; Bartlett and Silverman, 1989). In Xoo, both EPS and LPS are necessary for virulence. In our previous study (Rajeshwari and Sonti, 1997), SPVs were described to have no significant advantage over wild type in co-culture experiments and the same was obtained with the current set of SPVs (data not shown). Further studies are required to understand the reasons for the accumulation of EPS deficient phase variants in the prolonged stationary phase cultures of Xoo.

The two mechanisms observed in SPVs are transposition and SSM, which are established mechanisms of phase variation in bacteria. The *icaC* gene of the PIA EPS biosynthetic operon undergoes phase variation by *IS256* IS element transposition and SSM in *Staphylococcus aureus* (Kiem *et al*., 2004; Brooks and Jefferson, 2014). This suggests that bacteria have evolved more than one phase variation mechanism to regulate gene expression in the same locus. The *IS256* element might be involved in genome plasticity and adaptation to stress in *Staphylococcus aureus*. The introduction of *IS256* to a laboratory strain of *Staphylococcus aureus* lacking the *IS256* was shown to generate additional phenotype variants (Kleinert *et al*., 2017). The Xoo genomes have been described to accumulate IS elements both in diversity and copy number. IS elements have been attributed to contributing to genome rearrangements in the Xoo strain PXO99^A^, leading to genome plasticity and rapid evolution in Xoo strains (Salzberg *et al*., 2008). This study indicates the possibility that IS elements can also contribute to adaptations at a shorter timeline during the lifecycle of Xoo.

At the later stages of lesion development during Xoo infection, the rice leaves become dry and it is expected that this might lead to stresses such as nutrient limitation and desiccation. Thus it would be worthwhile to re-isolate bacteria from lesions of rice leaves inoculated with wild type Xoo strain to study if similar phase variants do occur during the *in-planta* growth of bacteria. Additional studies can also explore other phenotypes and growth conditions that would expand our knowledge about phase variation in plant pathogenic bacteria.

## Experimental procedures

### Bacterial strains, plasmids and growth conditions

Supplementary Tables S3 and S4 respectively list the bacterial strains and plasmids used in the study. Xoo strains were grown at 28°C in a 1% peptone-sucrose (PS) medium (Ray, Rajeshwari and Sonti, 2000). *E. coli* strains were grown at 37°C in Luria Bertani (LB) medium. The media was prepared with 1.5% Agar for plates. The antibiotics concentrations used were rifampicin (Rif) 50 µg/ml, kanamycin (Kan) 25 µg/ml, spectinomycin (Spec) 50 µg/ml, cephalexin (Ceph) 20 µg/ml, and cycloheximide (Cyclo) 80 µg/ml.

### Generation of stationary phase variants (SPVs)

SPVs were generated as previously described (Rajeshwari and Sonti, 1997). Individual mucoid colonies of wild type strain BXO43 were inoculated in 2 ml PS-Rif primary culture overnight. Each of the 2ml cultures were used as inoculum for 20 ml secondary culture in PS media and were grown for 48 hours. This was further maintained on a laboratory bench without shaking and was dilution plated on alternate days. Colonies were streaked out multiple times and screened for non-mucoid phenotype and one such colony exhibiting consistent phenotype in each inoculation was taken forward for characterisation.

### Non-mucoid phenotype complementation using genomic DNA library cosmids

SPV strains were transformed with *gum* operon containing cosmid pSD1 and LPS O-antigen biosynthetic gene cluster containing cosmid pSD5. A minimum of 16 transformed colonies selected on PSA-Kan plates per from each transformation were further streaked out for comparing non-mucoid phenotype complementation. Transforming the SPV strains with genomic library clones was performed by means of conjugation using S17-1 *E. coli* strain.

### LPS profiling on SDS-PAGE gels

LPS isolation was performed as previously described (Davis and Goldberg, 2012). Overnight cultures of Xoo strains were pelleted, washed, and resuspended in MilliQ. The cultures was washed and adjusted to OD_600_ = 1 in MilliQ water. From this resuspension 2 ml were pelleted and resuspended in 200 µl of 1X SDS-PAGE loading buffer and boiled for 15 minutes. To each sample, 200 µl of Tris saturated phenol was added, mixed and incubated at 65°C for 15 minutes and the aqueous layer was separated by adding 200 µl chloroform and centrifugation (). The hot-phenol extraction was repeated again and 100 µl of the aqueous layer was mixed with 200 µl of 2X SDS-PAGE loading dye and equal volumes were separated on 10% SDS-PAGE gels. The LPS bands were visualised by silver staining using ProteoSilver kit (Sigma-Aldrich, St. Louis, MO, USA).

### PCR screening for variations

The LPS O-antigen biosynthetic gene cluster of BXO1 (GenBank: AF337647) was compared with KACC 10331 sequence (GenBank: AE013598.1) for annotations. Thirteen overlapping primer pairs (listed in Supplementary Table S5) were designed manually for, LPS biosynthetic genes, *smtA* to *wxoD* for a PCR product of 500-600 bp and an overlap of ∼100 bp. Individual primer pairs were designed for genes transport proteins *wzt* and *wzm* and the *gum* operon gene *gumB*. The genomic DNA from Xoo strains were isolated and used as the template for PCR for each primer pair designed and PCR products were visualised on 1% agarose gels.

### Assessment of reversion phenotypes

Overnight cultures of Xoo strains were pelleted, washed in MiliQ water and adjusted to OD_600_ = 1. These suspensions were dilution plated to obtain single colonies. About 5 µl the bacterial suspensions for each strain were spotted on PS-Rif plates. The phenotypes were visually scored after 4-5 days of incubation.

### Virulence assay

Overnight culture of Xoo strains were pelleted, washed in MiliQ water and adjusted to OD_600_ = 1. Leaves of TN1 rice cultivar (40-60 days old) were clip inoculated by dipping surgical scissors in the bacterial suspension (Kauffman *et al*., 1973). The lesions were measured 15 days post inoculation.

### Isolation of bacteria from virulence lesions

About 3 cm of the region at the leading edge of the lesion was crushed in 1ml MilliQ water using a bead-beater and dilution plated on PS agar plates containing Rif, Ceph, and Cyclo to obtain single colonies. The mucoid phenotype of the colonies was scored after 5 days.

### Whole genome shotgun sequencing and data analysis

Whole genome shotgun (WGS) sequencing was performed for SPVs 1, 2, 3, 4, 5, 15, 22, 26, 31, 32, and 33. Fungal/Bacterial DNA MiniPrep kit (Zymo Research, Irvine, CA, USA) was used to extract the genomic DNA. Genomic DNA quality check was done using gel electrophoresis and NanoDrop 1000 instrument (Thermo Fisher Scientific, Waltham, MA, USA). Quantitative estimation of DNA was performed using Qubit 2.0 fluorometer (Invitrogen; Thermo Fisher Scientific). Nextera XT sample preparation kits (Illumina, Inc., San Diego, CA, USA) were used for Illumina paired-end sequencing libraries (2 × 250) with dual indexing adapters. Illumina sequencing libraries were sequenced on Illumina MiSeq platform (Illumina, Inc., San Diego, CA, USA). MiSeq control software was used to perform adapter trimming. Raw reads were assembled *de-novo* using spades v3.15.4. Annotation was done using prokka v1.01 (Seemann, 2014). The NCBI accession numbers for the WGS data are listed in Supplementary Table S6.

Genes related to LPS and EPS biosynthetic genes were fetched from Xoo, BXO1 from NCBI (CP033201). The status of these genes in the SPV isolates was checked at the protein level using tBLASTn and nucleotide level using BLASTn (Johnson *et al*., 2008). Genes not showing 100% identity or 100% coverage were further analysed by aligning using MEGA v.11.0.9 (Tamura, Stecher and Kumar, 2021) to look for non-synonymous mutations or indels. Further, SNPs were detected using ParSNP v1.2 (Treangen *et al*., 2014) taking BXO1 as a reference.

## Acknowledgements

V.N.M. acknowledges the University Grants Commission, Government of India for the Ph.D. fellowship. This work was supported by J. C. Bose fellowships to R.V.S. from the Science and Engineering Research Board, Government of India. R.V.S., P.B.P., and H.K.P. acknowledge CSIR, Government of India for various research project grants.

## Author contributions

VNM, HKP, and RVS, conceived the project and designed the experiments. VNM performed the experiments. KB, SK, and SM performed WGS and data analysis under the guidance of PBP. VNM, HKP, and RVS analysed the data and finalized the manuscript, which was approved by all the authors. PBP, HKP, and RVS contributed reagents/materials.

